# Plasticity and repeatability in spring migration and parturition dates with implications for annual reproductive success

**DOI:** 10.1101/2021.08.24.457438

**Authors:** Michel P. Laforge, Quinn M. R. Webber, Eric Vander Wal

## Abstract

Animals are faced with unprecedented challenges as environmental conditions change. Animals must display behavioral plasticity to acclimate to changing conditions, or phenotypic variation must exist within the population to allow for natural selection to change the distribution of trait values. The timing of migration and parturition relative to important annual environmental changes such as snowmelt and vegetation green-up and how they co-vary may influence reproductive success. We tested for plasticity and individual differences in migration and parturition timing as a function of the timing of snowmelt and green-up in a migratory herbivore (caribou; *Rangifer tarandus*, *n* = 92) using behavioral reaction norms. We tested whether timing of parturition, plasticity in parturition timing, or timing of green-up were correlated with calf survival. Migration and parturition timing were plastic to the timing of spring conditions, and we found moderate repeatability for migration timing, but no repeatability in timing of parturition. We detected a novel behavioral syndrome where timing of migration and timing of parturition were correlated. Our results suggest that observed shifts in caribou parturition timing in other populations are due to plasticity as opposed to an evolutionary response to changing conditions. We did not detect a correlation between annual reproductive success and either the timing of spring or plasticity to the timing of spring events. While this provides evidence that many populations may be buffered from the consequences of climate change via plasticity, we caution that a lack of repeatability in parturition timing could impede adaptation as climate warming increases.

**Significance Statement:** Animals have evolved to reproduce when resources are abundant. Climate change has altered the timing of annual events, resulting in earlier peaks in resource abundance. Animals can cope with this change in two ways. Individuals can display plasticity and alter the timing of reproductive activities to match the change in the environment, or consistent differences among individuals can result in sufficient variation to drive an evolutionary response. We tested these two alternative hypotheses in caribou (*Rangifer tarandus*) using an individual-based modelling framework. Caribou displayed plasticity in both when they migrated and gave birth, suggesting they can acclimate to changing conditions, but we did not find evidence of differences among individuals that would be likely to result in an evolutionary response.

---

Exhibiting life-history strategies to deal with fluctuations in habitat quality through time is fundamental for species living in seasonal environments (1). In many species, migration is viewed as a strategy to optimize use of seasonally available resources (2–4). In this context, migration will be most effective when individuals adjust their movement to match the phenology of their environment, which may serve as a cue to forecast spring conditions on summer range (5, 6). Optimizing opportunities to reproduce in seasonal environments also depends on individuals rearing young when resource availability is highest to properly finance reproduction (7, 8). Climate change, however, decouples migrants from optimally timing migration, and therefore reproduction. For example, warmer springs disproportionately affects the phenology, or timing of annual events, in lower trophic levels compared to higher trophic levels. The result is phenological asynchrony where consumers do not advance their phenologies to match that of their resource (9, 10). This asynchrony can lead to reduced survival and fitness, a phenological mismatch which can have significant population-level effects (10, 11). Plasticity in migratory and reproductive behavior in response to phenological shifts at lower trophic levels is relevant to species persistence as this plasticity likely buffers populations against adverse environmental change (12). Likewise, consistent among-individual differences can provide capacity for evolution within populations to adapt to changing conditions (13). Assessing potential outcomes of climate change on populations will necessitate elucidating how individuals are able to acclimate or adapt behaviors linked to annual reproductive success to inter-annual variation in resource phenology.

For terrestrial herbivores, spring migration is often driven by seasonal changes in the availability of high-quality forage resources (14) or snow cover that may presage future conditions on summer range (6). The forage maturation hypothesis suggests that herbivores should get the highest nutritional benefit by foraging on plants at an intermediate stage of growth when biomass is sufficient and digestibility is high (15, 16). Animals that forage on high-quality vegetation gain a nutritional benefit (17) and experience increased fat gain (18). While many populations or individuals track the emergence of high-quality forage as it matures along elevational and latitudinal gradients (14, 19), others use a strategy of “jumping” the green wave, arriving on summer range prior to when it peaks in forage quality (“green-up”; 19). Jumping the green wave may be related to snowmelt, with individuals tracking the progression of melting snow along migratory routes (6). The progression of snowmelt appears to be an important factor in migration timing for many northern ungulates. For example, in caribou (*Rangifer tarandus*) in Alaska and northern Canada, timing of arrival on summer range was correlated with the timing of snowmelt (20). Similarly for elk (*Cervus canadensis*) in Yellowstone National Park, departure from winter range was correlated with snowmelt date on winter range, and timing of arrival on summer range was associated with the timing of snowmelt on summer range and on migration routes (21). Using melting snow to time migrations may allow individuals to arrive on calving grounds to optimally take advantage of green-up during the calving season.

Animals that live in seasonal environments should be adapted to time their reproductive phenology such that the most energetically expensive times correspond with when resources are most highly abundant (7, 8). For ungulates, energetic requirements can more than double during the peak of lactation (22–24). In *Rangifer*, parturition date varies among populations as a function of the mean annual timing of green-up (25). Climate change disproportionately alters the phenology of lower trophic levels, resulting in phenological asynchrony where consumer breeding phenologies occur after the peak in resources at lower trophic levels, resulting in depressed reproductive success—a phenological mismatch (9, 26, 27). For example, changing sea ice phenology has altered vegetation phenology in Greenland, resulting in a phenological mismatch and reduced calf survival in caribou (28, 29). Long-term data suggests advancing migration and parturition dates in caribou over the last three decades, likely in response to changing resource phenologies (30) or spring temperatures (31). In the case of parturition, this is especially true for northern populations where climate warming has been more acute (32).

Populations can adjust their phenologies to cope with changes in the phenology of their resources in two ways. Individuals may acclimate their phenologies directly via behavioral plasticity. Alternatively, sufficient variation in phenotypes in the population might result in some individuals being better adapted to novel conditions (12, 33). If that variation is consistent among individuals, it may provide the pre-requisites for evolution. Plasticity is estimated over shorter, i.e., within generation, timescales by making repeated observations of individual phenologies and correlating them with environmental changes. Meanwhile, directly quantifying evolutionary responses requires data spanning multiple generations (34). The potential for evolution, however, can be inferred from shorter-term behavioral data. For example, natural selection requires traits that vary among individuals, and repeatability provides a measure of the proportion of variance in a trait that is attributable to differences among individuals (35). Highly repeatable behaviors are consistent within individuals but vary among individuals (36). Estimates of repeatability can therefore be used as tentative estimates for the potential for evolutionary responses when genetic data are unavailable (37)—see for example (38–40). Behavioral reaction norms (41) provide a framework to test whether contemporary changes in phenology are due to plasticity or a potential evolutionary response by modelling individual-level plasticity to changing environmental conditions while also estimating repeatability. Behavioral reaction norms can also quantify behavioral syndromes, the degree to which behaviors and behavioral plasticity are correlated (42). Exploring potential for syndromes in phenological traits, including migration and parturition timing, could demonstrate the importance of individuals properly timing migration to optimize the timing of parturition, with consequences to adaptation under changing environmental conditions.

We used behavioral reaction norm analyses of individual caribou (*n* = 92) in Newfoundland, Canada to test the non-mutually exclusive hypotheses that caribou exhibit plasticity in the timing of migration and parturition as a function of the timing of spring, and that these behaviors are repeatable traits providing variation that could result in an evolutionary response. We also tested for behavioral syndromes linking both timing of migration and parturition and their plasticity, and the phenological mismatch hypothesis by testing whether early springs reduced annual reproductive success. We tested several predictions, including:

1. Both timing of migration (P_1_a) and timing of parturition (P_1_b) would display plasticity as a function of changes in the annual timing of spring snowmelt. Individuals would both migrate early and give birth early in years where snowmelt occurred earlier. We also predicted that timing of parturition would display plasticity as a function of the timing of green-up (P_1_c), with individuals giving birth earlier when green-up was earlier.
2. Both timing of migration (P_2_a) and timing of parturition (P_2_b) would be repeatable behaviors. Individuals that migrate or give birth early do so consistently.
3. There is a correlation between timing of migration and timing of parturition (P_3_). Individuals that migrate early give birth early.
4. There is a correlation in degree of plasticity in migration timing and parturition timing (P_4_). Individuals that are more plastic in the timing of their migration are also more plastic in the timing of parturition.
5. Finally, we predicted that timing of green-up would influence calf survival, with lower survival in early springs which represent a phenological mismatch (P_5_).

## Results

We quantified the timing of spring migration and parturition of caribou as well as the timing of spring snowmelt and green-up (Figure 1). We fit two sets of behavioral reaction norm models: one with timing of migration and timing of parturition as response variables with timing of snowmelt as a fixed effect, and one with timing of parturition and calf survival to four weeks of age as response variables, with timing of green-up as a fixed effect. Both of the top behavioral reaction norm models included random slopes by individual ID for our response variables (timing of arrival on summer range, timing of parturition, and calf survival) as a function of our explanatory variables (timing of snowmelt and timing of green-up). This provided evidence for the importance of an individual × environment interaction (Δ DIC of both models to the next most supported model > 16). At the population-level, later snowmelt was significantly associated with later timing of arrival on summer range (β + 95% credible interval: 0.332 [0.192, 0.455], p < 0.001, supporting P_1_a) and nearly-significantly correlated with later parturition (0.138 [−0.006, 0.286], p = 0.067, partially supporting P_1_b), see Table 1 and Figure 2. We did not detect an effect of the timing of green-up on the timing of parturition (0.094 [−0.066, 0.270], p = 0.270, partial support for P_1_c), see Table 1 and Figure 2.

**Figure 1:**
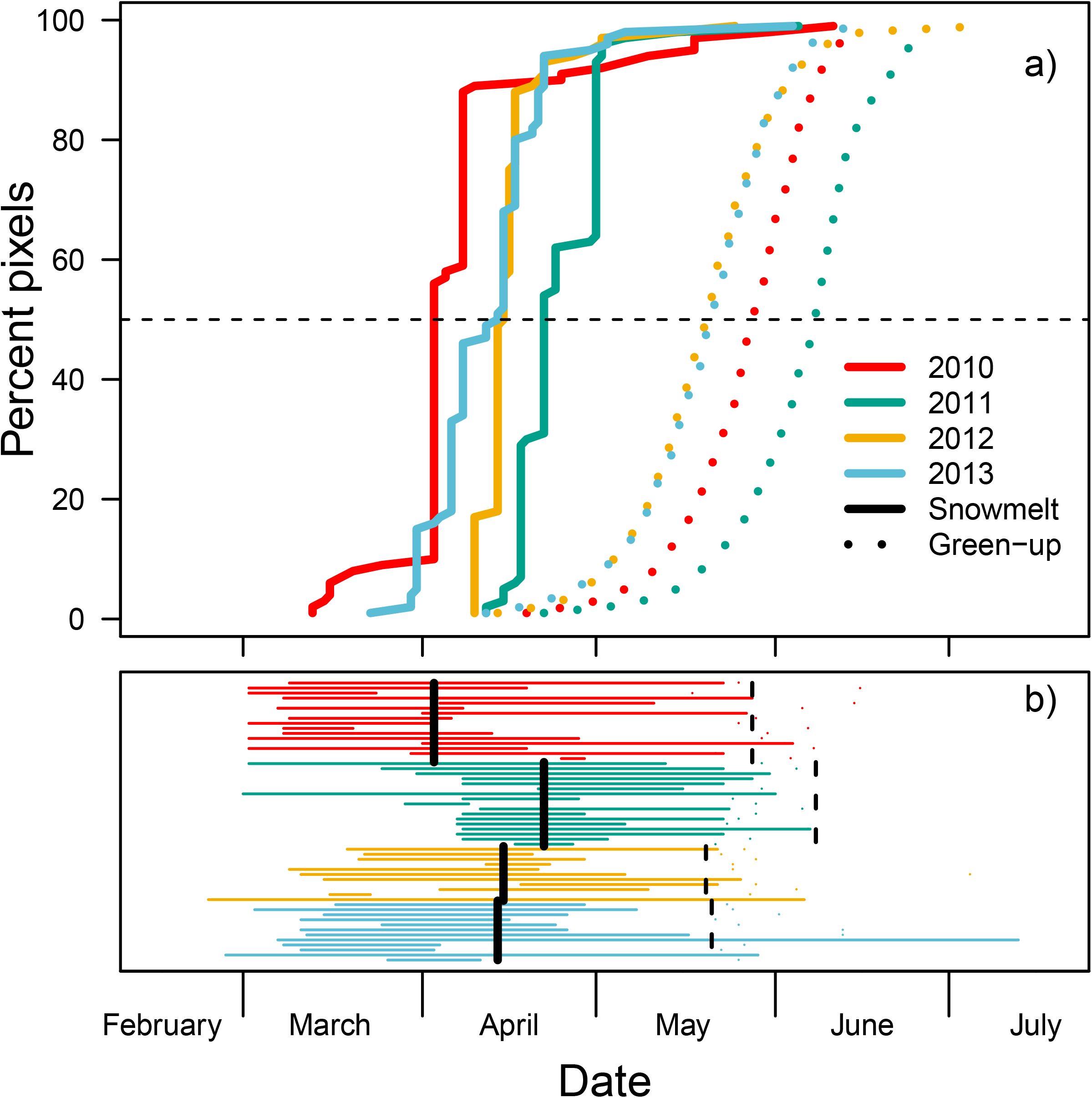
Phenology of snowmelt and green-up (a) and migration and parturition (b) for caribou (*Rangifer tarandus*, *n* = 32) in the Middle Ridge population, 2010–2013. a) represents the number of pixels within the population’s spring/summer range in which snow has melted (NDSI > 0, solid line) or that have reached the peak of green-up (instantaneous rate of green-up; dotted lines) in each spring. Colors represent different years, and the dashed line represents the median (the measure used for determining date of snowmelt/green-up). b) the timing of migration as horizontal lines for each individual in each year (each line’s extent represents the time they were migrating). Points represent the timing of parturition. Black vertical lines represent the median dates of snowmelt (solid) and green-up (dashed) for each year. See Figure S2 for plots using the other four populations used in this study.

**Figure 2:**
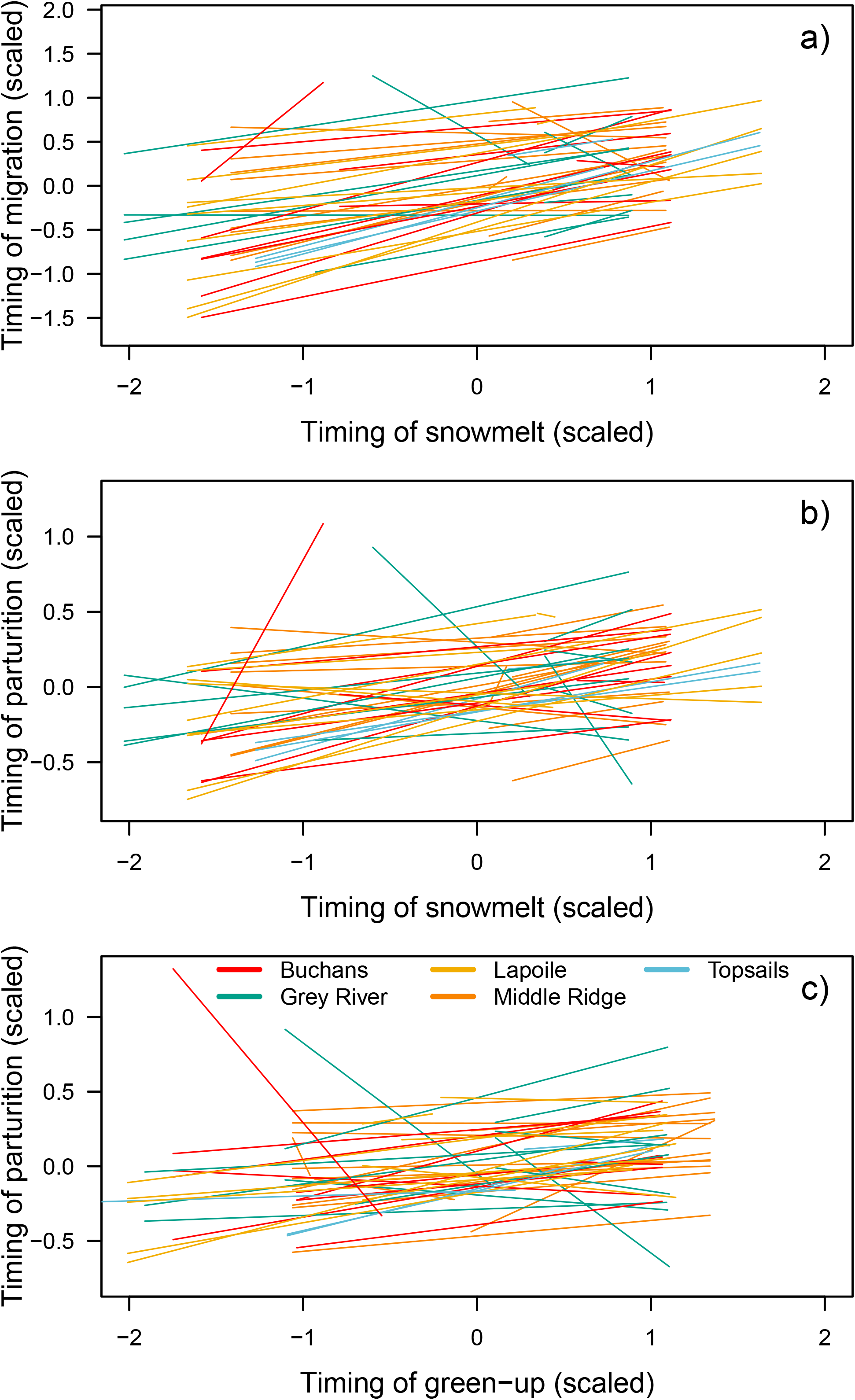
Mean-centred behavioral reaction norms for five caribou populations assessing timing of arrival on summer range and timing of parturition as a function of median date of spring snowmelt and green-up (see Materials and methods) for migratory caribou (*Rangifer tarandus*; *n* = 92) in Newfoundland, Canada. Each line represents a different individual. Panel a) represents timing of arrival on summer range as a function of timing of snowmelt, panel b) represents timing of parturition as a function of timing of snowmelt, and c) represents timing of parturition as a function of timing of green-up. Best linear unbiased predictors represent point estimates of the random effects from the mixed effects model.

**Table 1:** Estimates for fixed effects from the most parsimonious models from two bivariate Bayesian mixed effects models testing the effects of timing of snowmelt (model 1) and timing of green-up (model 2) on the timing of migration (model 1), timing of parturition (models 1 and 2) and probability of calf survival (model 2) for caribou (*Rangifer tarandus*, *n* = 92) in Newfoundland, Canada, 2007–2013. Estimates are presented with 95% credible intervals. The reference category for population is Buchans.

|  | Model 1: Snowmelt, arrival timing | Model 1: Snowmelt, parturition timing |
| --- | --- | --- |
| Arrival timing | - | 0.04 (-0.387, 0.457) |
| Parturition timing | 0.032 (-0.347, 0.367) | - |
| Snowmelt | 0.332 (0.192, 0.455) | 0.138 (-0.006, 0.286) |
| Grey River | -0.065 (-0.702, 0.559) | -0.021 (-0.532, 0.5) |
| Lapoile | -0.046 (-0.676, 0.471) | -0.002 (-0.469, 0.476) |
| Middle Ridge | -0.047 (-0.563, 0.472) | -0.024 (-0.473, 0.447) |
| Topsails | 0.09 (-0.568, 0.697) | -0.02 (-0.581, 0.535) |
|  | Model 2: Green-up, parturition timing | Model 2: Green-up, calf survival |
| Parturition timing | - | -0.037 (-0.375, 0.267) |
| Calf survival | 1.222 (-0.206, 2.817) | - |
| Green-up | 0.094 (-0.066, 0.27) | 0.316 (-0.377, 1.101) |
| Grey River | 0.063 (-0.401, 0.507) | -0.971 (-3.282, 1.091) |
| Lapoile | 0.099 (-0.38, 0.526) | -1.284 (-3.258, 0.665) |
| Middle Ridge | 0.027 (-0.395, 0.444) | 0.652 (-1.326, 3.132) |
| Topsails | 0.007 (-0.521, 0.554) | -0.968 (-3.34, 1.13) |

We found some evidence of among individual differences in the timing of migration across individuals, with repeatability for arrival on summer range being moderate (r [SD] = 0.377 [0.045], moderate support for P_2_a, Figure 3, red symbols). Overall repeatability for timing of parturition was quite low, suggesting that timing of parturition was not a trait that exhibited consistent individual differences in these populations (snowmelt model: 0.112 [0.004], green-up model: 0.051 [0.002] no support for P_2_b, Figure 3, blue-green symbols). We found evidence of a correlation between the intercept of migration timing and the intercept of parturition timing (0.679 [0.162, 0.986]), where early migrators also give birth earlier (P_3_; Figure 4a). We did not, however, find any evidence of a link between the plasticity in arrival on summer range and plasticity in timing of parturition (P_4_), individuals that were more plastic in the timing of their migration were not more plastic in the timing of parturition (−0.039 [−0.841, 0.757], Figure 4b). There was no support for our prediction that later green-up was correlated with higher calf survival (0.316 [−0.377, 1.101], p = 0.381, no support for P_5_, Table 1). We also did not find a significant correlation between timing of parturition and calf survival in an average environment (the mean date of green-up; −0.254 [−0.940, 0.568], Figure 4c). There was also no evidence that higher plasticity in parturition date resulted in higher overall calf survival (0.274 [−0.530, 0.951], Figure 4d).

**Figure 3:**
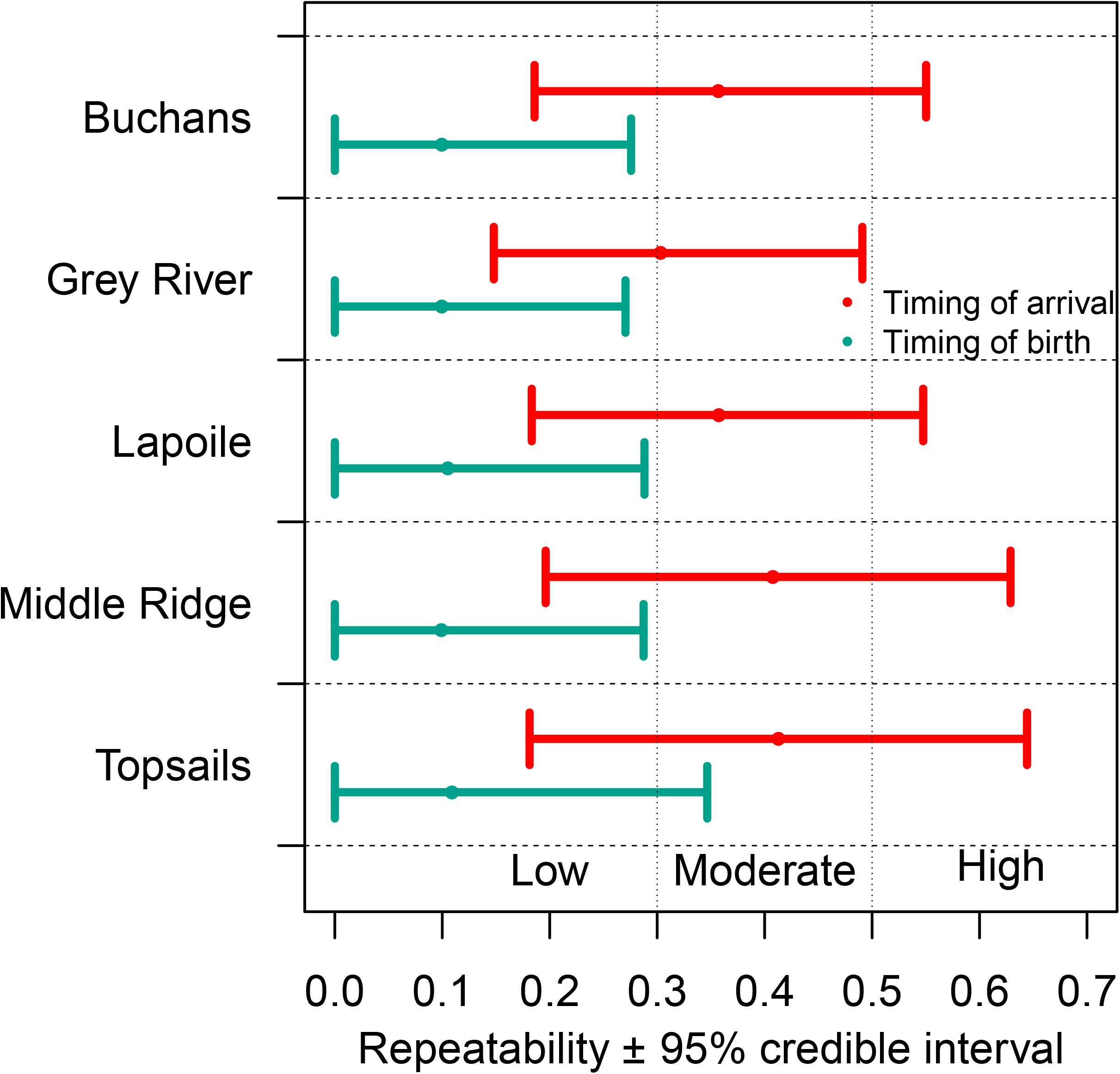
Repeatability estimates and 95% credible intervals for timing of arrival on summer range (red) and timing of birth (cyan) for caribou (*Rangifer tarandus*; *n* = 92) in Newfoundland, Canada. Repeatability estimates were derived from the top Bayesian mixed effects model describing timing of migration and parturition as a function of timing of snowmelt (the first model set).

**Figure 4:**
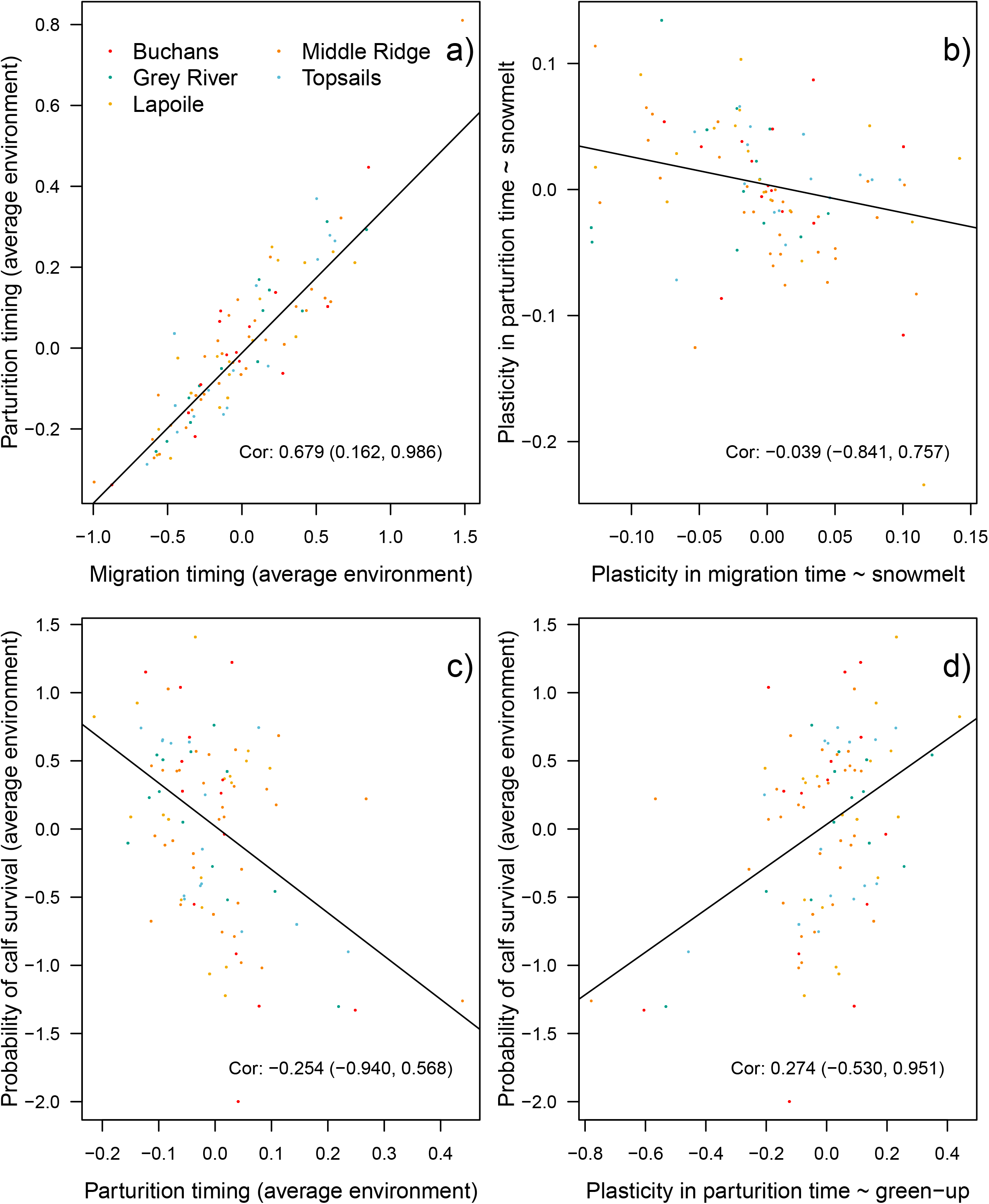
Correlations and 95% credible intervals between random slopes and intercepts from bivariate mixed effects behavioral reaction norm models quantifying timing of caribou (*Rangifer tarandus*; *n* = 92) migration, parturition, and calf survival as a function of the timing of spring snowmelt and green-up. Colors represent different populations. Panel a) represents the correlation between the intercept for timing of migration and the intercept for timing of parturition. Panel b) represents the correlation between the slopes (e.g., plasticity) of migration and parturition timing. Panel c) represents the correlation between the intercept for timing of parturition and the probability of calf survival to four weeks of age in an average environment. Panel d) represents the correlation between the plasticity in parturition time as a function of green-up and the probability of calf survival. Note, correlations are considered significant when credible intervals do not overlap zero and non-significant when credible intervals overlap zero.

## Discussion

An animals ability to match the phenology of life history events with changing snowmelt and green-up timing is vital for persistence, both across seasonal and inter-annual changes (8). We found evidence of individual differences in migration timing (P_2_a), such that some individuals consistently migrated earlier, and other individuals migrated later. Despite this, we found little evidence that timing of parturition was consistent among individuals (P_2_b). Our results suggest a primary role of plasticity in contributing to shifting life history phenology in caribou over the last few decades in other caribou populations (30, 32). With limited repeatability in parturition timing (r = 0.05–0.11), likely little of the variance is genetic (35), and therefore able to respond to selection. Migratory caribou in Newfoundland acclimated the timing of migration to the timing of snowmelt (P_1_a), and to a lesser extent, they also acclimated birth date to timing of snowmelt (P_1_b) but not to timing of green-up (P_1_c). This plasticity may have been sufficient to mitigate the effects of phenological mismatch on calving success in caribou, as we found no evidence for reduced annual reproductive success in early springs (P_5_). We also highlight a behavioral syndrome where individuals that migrate early also give birth early (P_3_). However, we did not find evidence that plasticity in timing of migration correlates with timing of parturition, suggesting that plasticity in these traits is independent (P_4_).

The results of our study indicate that migration timing is both a repeatable trait and plastic to the timing of snowmelt, outlining the importance of snow in many systems to synchronize animal phenology. In our populations, individuals migrated approximately one week prior to snowmelt, supporting the hypothesis that caribou use melting snow as a cue for when to migrate (6). Being plastic in migration timing to changes in snow phenology likely allows individuals to migrate at an optimal time to increase movement efficiency on ground or lakes that are still frozen (43) but without the impediment of deep snow. Moderate repeatability suggests that there are some individual differences in migration timing that could allow for evolutionary change as selective pressures on optimal migration dates change. Repeatability could have a genetic basis or a social one, for example, if individuals learn to migrate from conspecifics (44). The repeatability of arrival timing in our population (r = 0.377) was slightly higher than the mean value of arrival repeatability for long distance migratory birds (r = 0.31; 45). Consistent individual variation may be more favoured in environments where conditions on winter range are less reliable indicators of conditions on summer range, reducing the ability of individuals to be plastic to environmental variation (45). Relatively short-distance migrants like caribou in Newfoundland may be more adapted to migrate at the optimal time, reducing between-individual variation and therefore repeatability.

Caribou in our study displayed some plasticity in the timing of parturition to changes in the timing of snowmelt, suggesting individuals can acclimate their reproductive ecology to interannual environmental change. While timing of parturition is primarily correlated with the timing of the rut, environmental conditions have also been shown to impact the timing of parturition, including female body condition and spring temperature (31). Plasticity in breeding date is thought to buffer populations against the consequences of changes in optimal breeding date (46, 47). Earlier spring green-up has been shown to result in earlier birth in mule deer (*Odocoileus hemionus*; 48). Parturition date does vary across *Rangifer* population ranges to match local plant phenology (25, 32), and prior studies have documented long-term shifts in parturition date in caribou (30, 32). Repeatability of parturition date was low to moderate in red deer (*Cervus elaphus*, r = 0.19, 49) and even lower in our study on caribou (see Figure 3). Newfoundland has an unpredictable climate with large inter-annual variation driven primarily by the North Atlantic Oscillation (50, 51), which could have resulted in selection for plasticity in parturition date as opposed to selection for a specific optimal date (52, 53). Large variance in climate also suggests that trends of advancing parturition dates may be driven more by plasticity within generations as opposed to adaptive evolution across generations, especially given that these are still relatively short evolutionary timeframes.

It is unlikely that phenological mismatch is currently affecting caribou reproduction and fitness in Newfoundland. Timing of green-up did not significantly affect calf survival, although we note that this was due to large confidence intervals around a relatively large coefficient estimate. We cautiously suggest that early springs do not result in depressed reproductive success, likely because this scenario is mitigated by plasticity in the timing of parturition allowing individuals to avoid significant mismatch. Despite indications of phenological mismatch in caribou (29), other studies have failed to detect a significant effect of spring asynchrony on caribou forage availability (54). Some of our results however raise the spectre that caribou may not be able to continue to acclimate or adapt to future conditions. If individuals are plastic to a cue that is increasingly unreliably correlated with fitness-maximizing resources, this can result in phenological mismatch despite plasticity. For example, differences in the amount of warming experienced in the Netherlands in early versus late spring resulted in mismatch between great tits (*Parus major*) and their caterpillar prey, as the birds displayed plasticity to the former and the caterpillars the latter (55, 56). Reproductive success in caribou appears to be linked to timing parturition to coincide with peak green-up (29); however, we found that individuals were more plastic in the timing of birth in response to the timing of snowmelt than to green-up itself. The timing of green-up was only loosely correlated with the timing of snowmelt (R^2^ = 0.21), with green-up only advancing by 0.30 days for each day that snowmelt advanced (Figure S3). It is likely that climate change will continue to disrupt the predictability of the relationship between these two phenological events, which may result in caribou using an increasingly unreliable cue to time reproduction.

Evidence from our study suggests that earlier parturition date in our populations and potentially in other *Rangifer* populations (32) is primarily driven by plastic changes as individuals acclimate to changing conditions as opposed to natural selection acting to select for individuals that give birth earlier in the season. Repeatability is often considered to represent an upper bound to heritability (35), and as such the low repeatability of parturition date we observed suggests it is unlikely that parturition date is heritable in caribou. Plasticity alone, without any evolutionary change, could simply delay population decline if populations are driven beyond the limits of their ability to acclimate (57, 58). Experimental evidence from both frogs and passerines suggest that current rates of evolution are insufficient to account for observed shifts in reproductive phenologies (59, 60).

Our behavioral reaction norm framework allowed us to test for correlations in individual responses in environmental change to timing of snowmelt and green-up. For both of our model sets, the top model included random slopes for individuals, suggesting the existence of among-individual variation in the degree of plasticity to environmental change. However, we only detected significant among-individual correlations among two traits: timing of migration and timing of parturition, providing evidence of a link between these two behavioral traits. Counter to our fourth prediction however, we did not find a link between plasticity in timing of migration and plasticity in timing of parturition, suggesting that the ability to acclimate migration timing is not related to plasticity in parturition date. We also failed to detect a significant relationship between plasticity in parturition date and reproductive success. For a telemetry study, we have a large sample of individuals, repeated across several years, and replicated among herds. Despite our relatively large sample size, bivariate mixed effects models, like those used in this study, are data hungry (61). While some of our conclusions may be based on what could be considered modest statistical inference, we suggest that finding strong inference in our data will be more challenging than in contexts where behavioral tests can be performed across a range of environments that can be replicated numerous times in a single year to obtain a robust sample size (i.e, 62, 63).

Climate change represents an imminent threat to migratory animals such as caribou (4, 64). Despite indications that phenologies of some migratory species are shifting to compensate for changes at lower trophic levels (30, 32), understanding the mechanism remains an important question. By empirically demonstrating individual plasticity to changing environmental conditions and low repeatability in parturition time, our results suggest that much of the observed contemporary shifts in phenology are due to plastic responses to environmental change as opposed to an evolutionary response. Plasticity also appeared to buffer populations against depressed fitness, as we found no effect of the timing of spring on annual reproductive success. We demonstrate an important link between migration timing and parturition timing, suggesting that phenological synchrony is a two-part process in caribou—individuals must not only give birth when conditions are optimal, but this synchrony is also dependent on migration timing. Caribou are globally threatened (65), and as a northern ungulate exposed to high levels of climate change-induced warming, provide an important litmus test for understanding the impact of climate change on migratory herbivores globally.

## Methods

### Study site

We conducted our study on the island of Newfoundland, Canada (~47° 44’ N, 52° 38’ W to 51° 44’ N, 59° 28’ W). Caribou habitat in Newfoundland primarily consists of coniferous forest and mixed wood forests dominated by balsam fir (*Abies balsamea*), black spruce (*Picea mariana*), and white birch (*Betula papyrifera*), interspersed with bog and heath habitats. Barren rock and lakes are also common features in Newfoundland (see study area map, Figure S1).

### Data collection and cleaning

GPS collars (Lotek Wireless Inc., Newmarket, ON, Canada, GPS4400M collars, 1,250 g) were deployed on 112 adult female caribou from five populations on Newfoundland between 2007– 2013. Caribou were captured by darting from a helicopter. GPS fixes were obtained every 1–5 hours depending on season and collar. All animal capture and handling followed guidelines from the Canadian Council on Animal Care. We initially had data from 112 individuals over 326 ID-years. Cleaning the data and removing ID-years with insufficient data, and non-parturient individuals left us with data for 103 individuals across 294 ID-years. A further nine individuals and 78 ID-years were dropped after removing individuals that did not migrate at least 30 km. Our final dataset consisted of data for 92 caribou across 212 ID-years.

### Defining timing of snowmelt and green-up

We used two measures of phenological change to quantify the timing of spring events for caribou populations in Newfoundland, the timing of snowmelt and the timing of green-up. We used the timing of snowmelt as the presumed driver of spring migration timing as snowmelt has been shown to correspond to the timing of migration of caribou in Newfoundland (Laforge et al 2021). We used the normalized difference snow index (NDSI) derived from daily MODIS data at a spatial resolution of 250 × 250 m to determine the date of snowmelt. We determined the day that each pixel first had a recorded negative value of NDSI as the date upon which each pixel was considered snow-free. We used the timing of snowmelt within each population’s range (99% MCP of all locations from the start of spring migration to three weeks after parturition) to define the timing of snowmelt for each individual in each year as the median date of when pixels in the population-range became snow-free.

To quantify the timing of plant green-up, we used the instantaneous rate of green-up (IRG). IRG represents the first derivative of a series of normalised difference vegetation index values at a given location. We used data from both MODIS satellites (Terra and Aqua), each of which produced 16-day composite NDVI images at a resolution of 250 × 250 m. The sensors on each of the two satellites produce composite images at opposite times (i.e., phased), so combining the two data records provides an 8-day temporal resolution. We first set to NA any NDVI values where the snow cover band of the MODIS data indicated snow cover and replaced it with the 3^rd^ percentile of all snow-free observations at that pixel. This procedure ensures that the resulting curves are only plotting the change in plant growth and not confounded by melting snow (14, 19). We also set any pixels contaminated by cloud (~10.0%) to NA. For each location in our study area and for each year of our study, we used a 3-observation moving median filter to smooth the time series then fit a logistic curve to the time series of NDVI values at that location. We then calculated as IRG the first derivative of this curve and determined the date that IRG had the highest value. This date represented the day in which plant growth was occurring fastest and therefore represented the highest nutritional quality for caribou (17, 18). To calculate the date of green-up in each herd’s range each year, we generated a population-level MCP range (as defined above for snowmelt) and calculated the median date each year that pixels in the home range reached peak IRG. The median date of snowmelt across ranges and years was Apr 27, and the median date of green-up was May 29 (see Figure 1).

### Defining caribou migration timing

To define timing of migration, we used Migration Mapper V. 2.0 (66). Migration Mapper plots GPS locations on a map along with a profile of net squared displacement to allow the user to visually inspect these profiles to specify when individuals begin and complete migratory movements. As we were only interested in spring migration, we only quantified the timing of departure from winter range and arrival on summer range. We used the date of arrival on summer range as our measure of timing of migration. We removed any individuals that did not migrate at least 30 km. The median date for departure from winter range was Mar 25 (range: Feb 9–May 19). The median duration of migration was 43 days (range: 4–128) covering a median distance of 62.2 km (range: 30.5–174.9). The median date for arrival on summer range was May 10 (range: Mar 20–Jul 13, see Table 1).

### Quantifying timing of parturition and annual reproductive success

We used the method developed by DeMars et al. (67) and validated by Bonar et al (68) to define calving date and whether calves survived to four weeks of age. This method quantifies birthing events by detecting constraints on movement in females who must stop to give birth and whose movements are constrained by calves-at-heel that have a slower movement rate. Females whose calves die in the first four weeks of life display a sudden return to baseline movement rates, whereas females whose calves display a gradual return to baseline rates as calves are able keep pace with their mothers. We used a population-based method to detect calving and calf mortality events that examined three-day average movement rates of collared females to ascertain whether females gave birth that year and quantify the date of parturition. In some instances, the model output suggested that individuals gave birth in the first day of the time series provided to the model. In these instances, we manually validated parturition dates by inspecting a plot of daily movement rates. These data were validated using the Middle Ridge population, in which the method correctly classified 100% of parturition events (68). The median date of calving was May 30 (range: May 18–Jul 13), and 60% of calves survived to four weeks of age (see Table S1).

### Behavioral reaction norms

Behavioral reaction norms quantify how individual behavior changes across an environmental gradient. To evaluate how individual phenotypes for timing of migration, parturition, and annual reproductive success are expressed across an environmental gradient (timing of snowmelt or green-up), we quantified behavioral reaction norms (BRNs; 41) using two sets of bivariate Bayesian mixed effects models (R package *MCMCglmm*, version 2.29; 61, 69). We predicted that timing of migration and parturition would both be affected by timing of snowmelt, and that timing of parturition would also be affected by the timing of green-up, as would calf survival. Therefore, the response variables for our first set of models were the date of arrival on summer range and the timing of parturition, with timing of snowmelt used as our main explanatory variable. For our second model, response variables were timing of parturition and whether caribou calves survived to four weeks of age, with timing of green-up used as the explanatory variable. Variables were scaled independently for each population. Models were fit with uninformative priors (34) and Gaussian error structures for timing of migration and parturition, and with a categorical (binomial) error structure for calf survival. We ran models with a total of 420,000 iterations with a burn-in of 20,000 and a thinning rate of 100. We evaluated eight models using different combinations of random and fixed effects structures (Table S2). We tested for the effect of individual ID, and for the effect of an individual × environment interaction (i.e., an interaction between individual caribou and relative date of snowmelt or green up; 70) in the random terms. In our BRN analyses, fixed effects were used to control for changes in the random effects. In each of our two model sets, we chose the model with the lowest deviance information criterion (DIC, see Table 1). We extracted best linear unbiased predictors (e.g., random intercept and slope estimates for each ID-year) and calculated repeatability (*r*) of BRN intercepts for migration date, parturition date, and calf survival as the amount of between-individual variance (*V_ind_*) attributable to the residual variance among groups (*V_res_*) for each trait (71):

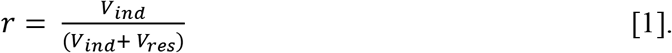

We examined the correlation between the slope and intercept of best linear unbiased estimators (random effects) for both variables in each model to examine the relationships between individual differences and plasticity within and between the traits (61; Table S3).

## Acknowledgements

The authors would like to acknowledge members of the Newfoundland and Labrador Wildlife Division including S. Moores, B. Adams, C. Doucet, W. Barney, F. Norman, R. Otto, J. Neville, P. Saunders, T. Porter, P. Tremblett, S. Gullage, T. Hodder, D. Jennings, and J. McGinn for managing and curating data on Newfoundland caribou. We thank T. Bergerud and S. Mahoney for their vision in initiating much of the work on caribou in Newfoundland. Parturition and survival data were calculated by M. Bonar. The authors would like to acknowledge J. A. Merkle, K. P. Lewis and S. J. Leroux for valuable comments on previous versions of this manuscript, as well as feedback from J. A. Aubin, J. Balluffi-Fry, M. Bonar, K. Kingdon, L. Newediuk, C. M. Prokopenko, A. L. Robitaille, and S. Zabihi-Seissan. Funding was provided by Natural Sciences and Engineering Research Council of Canada grants to M. Laforge (PGS-D), Q. Webber (Vanier CGS), and E. Vander Wal (Discovery). M. Laforge was also supported by Memorial University of Newfoundland’s A.G. Hatcher Memorial Scholarship.

