## Supplementary Materials for "Plasticity and repeatability in spring migration and parturition dates with implications for annual reproductive success"

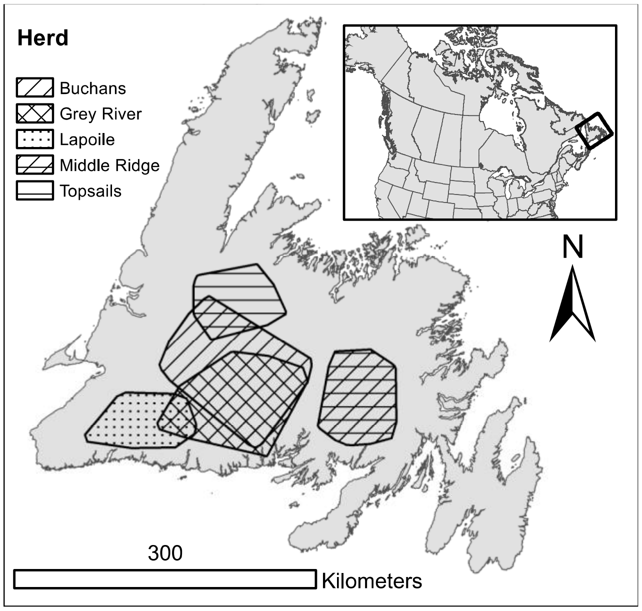


**Figure S1** The island of Newfoundland with the location of the five caribou (*Rangifer tarandus*) populations used in this study.

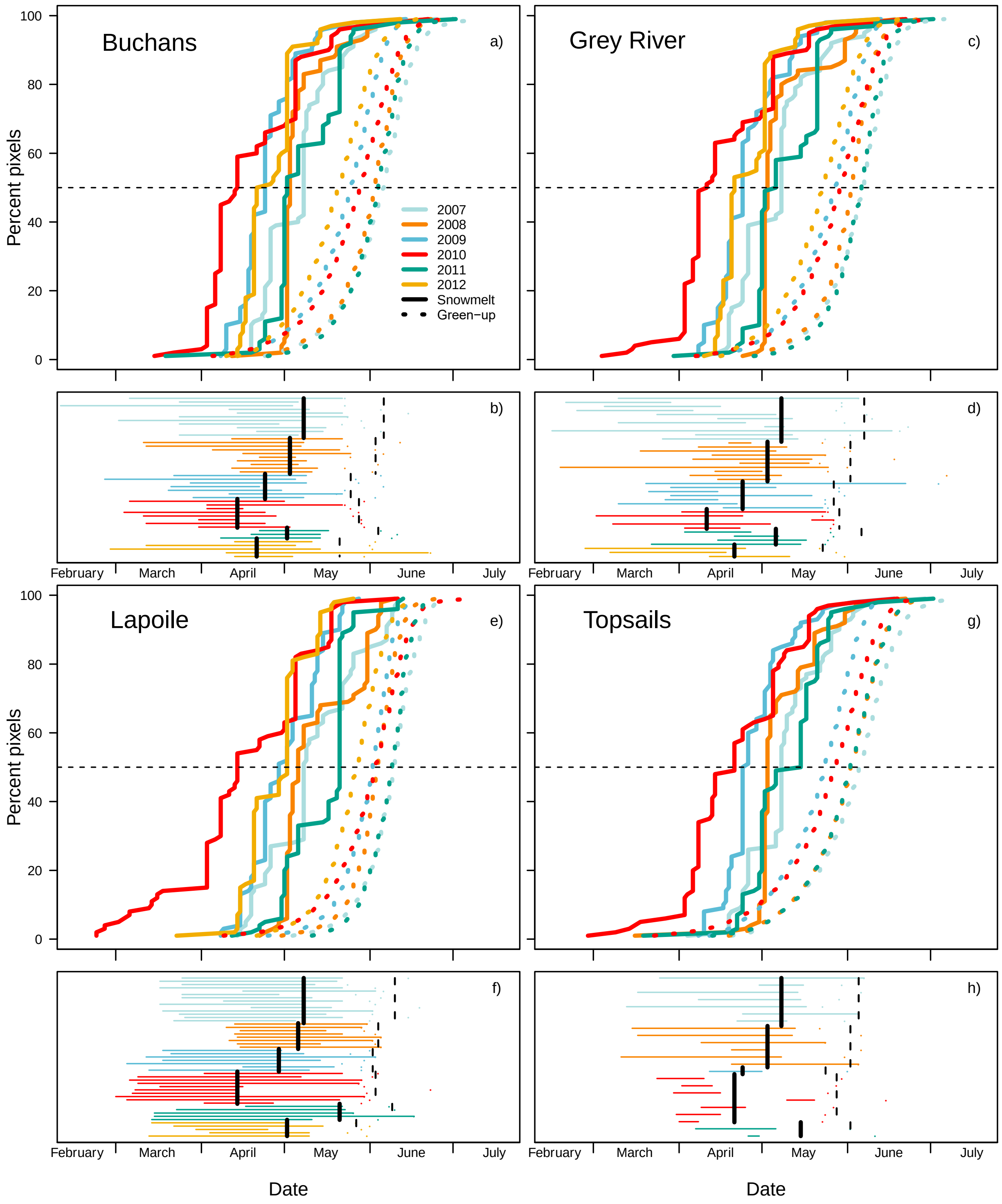


**Figure S2** Phenology of snowmelt and green-up and migration and parturition for caribou (*Rangifer tarandus*, $n$ = 60) in the Buchans (a, b), Grey River (c, d), Lapoile (e, f), and Topsails (g, h) populations, 2007–2012 (no data for Topsails, 2012). a), c), e), and g) represents the number of pixels within the population's spring/summer range in which snow has melted (NDSI > 0, solid lines) or that have reached the peak of green-up (instantaneous rate of green-up; dotted lines) in each spring. Colors represent different years, and the vertical dashed lines represent the median (the measure used for determining date of snowmelt/green-up). b), d), f), and h) represent the timing of migration as horizontal lines for each individual in each year (each line's extent represents the time they were migrating). Points represent the timing of parturition. Black vertical lines represent the median dates of snowmelt (solid) and green-up (dashed) for each year. See Figure 1, main text for plots for the Middle Ridge herd.

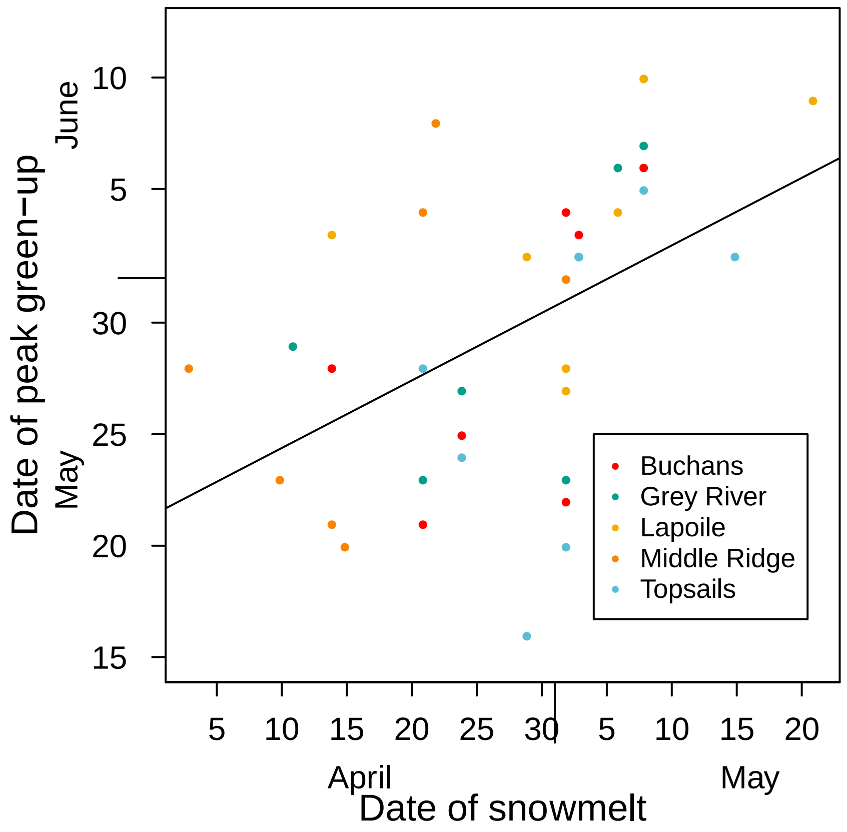


**Figure S3** Relationship between date of snowmelt and date of green-up in the ranges of five caribou (*Rangifer tarandus*) populations in Newfoundland, 2007–2013. There was a significant relationship between green-up date and snowmelt date, however later snowmelt only resulted in green-up occurring 0.303 (95% CI: 0.114, 0.492) days later (adjusted $R^2 = 0.21$).

**Table S1** Summary statistics of caribou (*Rangifer tarandus*, $n$ = 92) sample size and spring migration timing, duration, and distance across five populations in Newfoundland, Canada from 2007–2013.

| | All Populations | Buchans | Grey River | Lapoile | Middle Ridge | Topsails |
|---|---|---|---|---|---|---|
| Number of individuals | 92 | 14 | 13 | 18 | 32 | 15 |
| Number of ID years | 212 | 44 | 40 | 49 | 56 | 23 |
| median + [range], start date of migration | Mar 25 [Feb 09, May 19] | Mar 31 [Feb 09, Apr 22] | Mar 29 [Feb 14, May 19] | Mar 22 [Mar 01, Apr 19] | Mar 18 [Feb 23, Apr 25] | Apr 07 [Mar 11, May 10] |
| median + [range], end date of migration | May 10 [Mar 20, Jul 13] | May 08 [Apr 14, Jun 22] | May 07 [Mar 29, Jun 22] | May 18 [Apr 13, Jun 17] | May 04 [Mar 20, Jul 13] | May 10 [Apr 08, Jun 07] |
| median + [range], duration of migration (days) | 43 [4, 128] | 40.5 [13, 84] | 39.5 [7, 123] | 51 [25, 94] | 43 [4, 128] | 19 [4, 74] |
| median + [range], distance of migration (km) | 62.2 [30.5, 174.9] | 105.3 [42, 143] | 58 [36.4, 174.9] | 99.8 [35.9, 173.5] | 48.9 [30.5, 82.3] | 48 [32.1, 139.1] |
| median + [range], date of calving | May 30 [May 18, Jul 13] | May 31 [May 20, Jun 23] | May 28 [May 22, Jul 07] | Jun 01 [May 22, Jun 23] | May 29 [May 18, Jul 13] | May 31 [May 22, Jun 15] |
| Proportion calves survive | 0.6 | 0.66 | 0.55 | 0.51 | 0.7 | 0.52 |

| | | | | | | |
|---|---|---|---|---|---|---|
| Median + [5th and 95th percentile] date of snowmelt | Apr 27 [Apr 12, May 19] | Apr 27 [Apr 16, May 21] | Apr 27 [Apr 15, May 24] | May 03 [Apr 09, May 25] | Apr 16 [Apr 08, May 03] | May 01 [Apr 13, May 23] |
| Median + [5th and 95th percentile] date of green-up | May 29 [May 04, Jun 13] | May 28 [May 01, Jun 13] | May 30 [May 04, Jun 13] | Jun 03 [May 13, Jun 15] | May 28 [May 06, Jun 12] | May 27 [Apr 26, Jun 13] |

**Table S2** Random and fixed effect structures and Δ deviance information criterion (DIC) for candidate behavioral reaction norm models for migratory caribou (*n* = 92) in Newfoundland, Canada. Two model sets are compared: one set with timing of migration (arrival on summer range) and timing of parturition as co-response variables and timing of snowmelt as the explanatory variable, and one set with timing of parturition and calf survival as co-response variables with timing of spring green-up as the explanatory variable. Timing of snowmelt/green-up is the median date in which pixels in the population's range became snow-free (migration and parturition models) or reached peak green-up (parturition and calf survival models) that spring.

| Model | Fixed effects | Random effects | **Δ** DIC migration and parturition | **Δ** DIC parturition and calf survival |
|---|---|---|---|---|
| Null | Population | None | 81.2 | 50.4 |
| Population differences, no plasticity | Population | Population | 81.2 | 60.8 |
| Individual differences, no plasticity | Population | Individual | 56.0 | 35.9 |
| Overall plasticity, no population/individual differences | Population + Timing of snowmelt/green-up | None | 58.4 | 50.0 |

| | | | | |
|---|---|---|---|---|
| Overall plasticity, population differences | Population + Timing of snowmelt/green-up[1] | Population | 58.5 | 50.1 |
| Overall plasticity, individual differences | Population + Timing of snowmelt/green-up | Individual | 16.0 | 34.4 |
| Random slopes, population × environment interaction | Population + Timing of snowmelt/green-up | Population × Timing of snowmelt/green-up | 59.6 | 37.6 |
| Random slopes, individual × environment interaction | Population + Timing of snowmelt/green-up | Individual × Timing of snowmelt/green-up | 0 | 0 |

**Table S3** Slope and intercept correlations (plus upper and lower 95% credible interval) between timing of arrival on summer range, timing of birth, and annual reproductive success for caribou (*Rangifer tarandus*, n = 92) in Newfoundland Canada. Results from the top Markov chain Monte Carlo bivariate generalized linear regression models, along with an interpretation of what each correlation suggests.

| Traits | Interpretation | Estimate | Lower | Upper |
|---|---|---|---|---|
| Migration time intercept, birth time intercept | Timing of migration (arrival on summer range) and timing of parturition in an average environment are correlated. Individuals that migrate earlier give birth earlier (see Figure 3a, main text). | 0.68 | 0.16 | 0.99 |
| Migration time intercept, migration time slope | No strong correlation between timing of migration in an average environment and amount of plasticity in migration timing. | -0.36 | -0.97 | 0.47 |
| Birth time intercept, birth time slope | No correlation between timing of birth in an average environment and plasticity in timing of birth. | 0.19 | -0.76 | 0.91 |
| Migration time slope, birth time slope | No correlation between plasticity in migration timing | -0.04 | -0.84 | 0.76 |

| | | | | |
|---|---|---|---|---|
| | and parturition timing (see Figure 3b, main text). | | | |
| Migration time slope, birth time intercept | No correlation between plasticity in migration timing and timing of birth in an average environment. | -0.21 | -0.92 | 0.66 |
| Migration time intercept, birth time slope | No strong correlation between timing of migration in an average environment and plasticity in timing of parturition. | 0.27 | -0.68 | 0.92 |
| Calf survival intercept, birth time intercept | No strong correlation between probability of calf survival and timing of parturition in an average environment (see Figure 3c, main text). | -0.25 | -0.94 | 0.57 |
| Calf survival intercept, calf survival slope | No strong correlation between average probability of survival and whether probability of survival changes as a function of green-up. | -0.26 | -0.90 | 0.54 |
| Birth time intercept, birth time slope | Potentially negative correlation suggests that individuals that give birth late may be less plastic in the timing of parturition. | -0.52 | -0.99 | 0.42 |

| | | | | |
|---|---|---|---|---|
| Calf survival slope, birth time slope | Likely negative correlation suggests that increased plasticity in parturition time results in less heterogeneity in annual reproductive success across a gradient of green-up dates. | -0.57 | -0.97 | 0.10 |
| Calf survival slope, birth time intercept | Potentially positive correlation suggests individuals who give birth later are more likely to have their calf survive when green-up is later. | 0.49 | -0.41 | 0.98 |
| Birth time slope, calf survival intercept | Potentially positive correlation suggests that greater plasticity in parturition time may increase reproductive success (see Figure 3d, main text). | 0.27 | -0.53 | 0.95 |
